# The use of negative control outcomes in Mendelian Randomisation to detect potential population stratification or selection bias

**DOI:** 10.1101/2020.06.01.128264

**Authors:** Eleanor Sanderson, Tom G Richardson, Gibran Hemani, George Davey Smith

## Abstract

A key assumption of Mendelian randomisation (MR) analysis is that there is no association between the genetic variants used as instruments and the outcome other than through the exposure of interest. Two ways in which this assumption can be violated are through population stratification and selection bias which can introduce confounding of the relationship between the genetic variants and the outcome and so induce an association between them. Negative control outcomes are increasingly used to detect unobserved confounding in observational epidemiological studies. Here we consider the use of negative control outcomes in MR studies. As a negative control outcome in an MR study we propose the use of phenotypes which are determined before the exposure and outcome but which are likely to be subject to the same confounding as the exposure or outcome of interest. We illustrate our method with a two-sample MR analysis of a preselected set of exposures on self-reported tanning ability and hair colour. Our results show that, of the 33 exposures considered, GWAS studies of adiposity and education related traits are likely to be subject to population stratification and/or selection bias that is not controlled for through adjustment and so any MR study including these traits may be subject to bias that cannot be identified through standard pleiotropy robust methods.

## Introduction

When the observed association between an exposure, *X*, and an outcome, *Y*, is confounded by an unobserved variable conventional regression analysis will produce misleading estimates of the effect of the exposure on the outcome. If genetic variants – usually single nucleotide polymorphisms (SNPs) - are available which reliably predict the exposure variable but do not have an effect on the outcome through any other pathway, then they are valid instrumental variables (IVs) and can be used in a Mendelian randomization (MR) analysis to obtain unconfounded evidence of the effect of the exposure on the outcome.(1, 2) A key assumption for MR to give consistent estimates of the causal effect of an exposure on the outcome is that the SNPs used as instruments are not associated with the outcome other than through the exposure.(3) One way in which this assumption may be violated is through structure in the genetic variants in the population studied affecting both the exposure and the outcome. Structure of this type could arise through population stratification in the exposure and outcome variables or selection bias affecting whether individuals with particular characteristics are more or less likely to be involved in the study.(4, 5)

MR analyses are often conducted by comparing summary data estimates of SNP-exposure and SNP-outcome associations gleaned from two independent but homogeneous study populations. This is referred to as two-sample summary data MR.(6) For the MR estimate of the causal effect of the exposure on the outcome to be a consistent estimate of the effect of the exposure on the outcome the genetic variants must satisfy the following assumptions.

IV1: the variants must be associated with the exposure *X* (the “relevance” assumption);
IV2: the variants must be independent of all (observed or unobserved) confounders of *X* and *Y*, as represented by U (the “exchangeability” assumption); and
IV3: the variants must be independent of the outcome *Y* given the exposure *X*, (the “exclusion restriction”).

These assumptions are illustrated in Fig. 1 and are explained in detail elsewhere.(3, 6, 7) Two common causes of confounding of the instrument and outcome in MR, violating IV2, are population stratification and selection bias.

**Figure 1:**
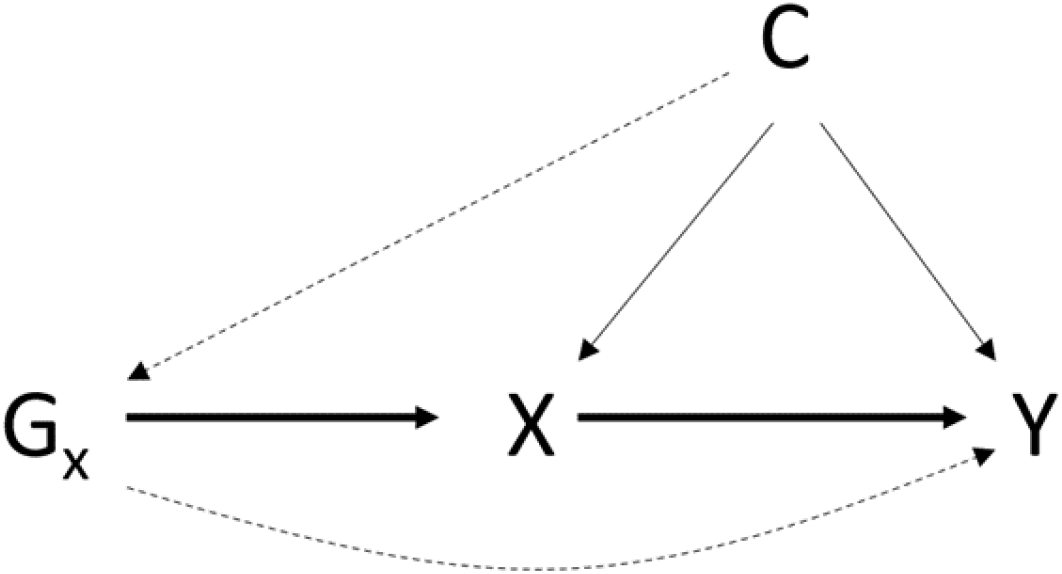
Instrumental variable assumptions. X is the exposure of interest, Y is the outcome of interest, G_X_ are the genetic variants associated with X used as instruments, C is a confounder of the exposure outcome relationship. Assumption IV1 is illustrated by the bold line from G_X_ to X. Violations of assumptions IV2 an IV3 are given by the dashed lines from C to G_X_ and from G_X_ to Y respectively.

Population stratification, where different sub-populations within the population being studied have different allele frequencies and also have different distributions of the phenotype will confound the results from the GWAS and lead to potentially spurious or inflated association between SNPs and the phenotype which are due to the structure of the population and not due to a direct effect of the SNP on the phenotype.(2, 8) Within GWAS studies population stratification is often controlled for by adjusting for the top principal components from a principal components analysis of the genetic variants.(9) Recently it has become increasingly popular to undertake GWAS using linear mixed models which allows them to account for genetic confounding of common variants more accurately, and improve power, by jointly modelling the contribution of all measured variants.(10–12) Generally MR analyses have assumed that any population level genetic structure is fully accounted for in the GWAS study that gives the estimates of the effects of the SNPs on the phenotypes for the analysis. There is increasing evidence however that this assumption does not hold for a number of phenotypes, particularly for social phenotypes, such as educational attainment, that may vary geographically.(4, 5) Haworth et al (2019) show that genetic variants are associated with a number of variables including location of birth in UK Biobank and this association cannot be fully accounted for by standard principal components analysis.(4) Population stratification introduces bias in MR studies by creating an association between the SNPs used as instruments and the outcome illustrated in Fig. 2. Therefore, any MR analyses based on the results from a GWAS study will potentially be biased if that GWAS does not fully account for any ancestral population structure that could lead to population stratification.(5, 13) This bias is likely to be largest when the outcome phenotype in a MR study is subject to population stratification that has not been fully accounted for. However it will also bias effect estimates in a two-sample MR analysis when the exposure phenotype is subject to population stratification by causing the estimated association between the SNP and the exposure to be mis-specified. As well as increasing or decreasing the size of the observed association this bias could create an apparent causal association between the exposure and the outcome when none exists, or mask a true association.

**Figure 2:**
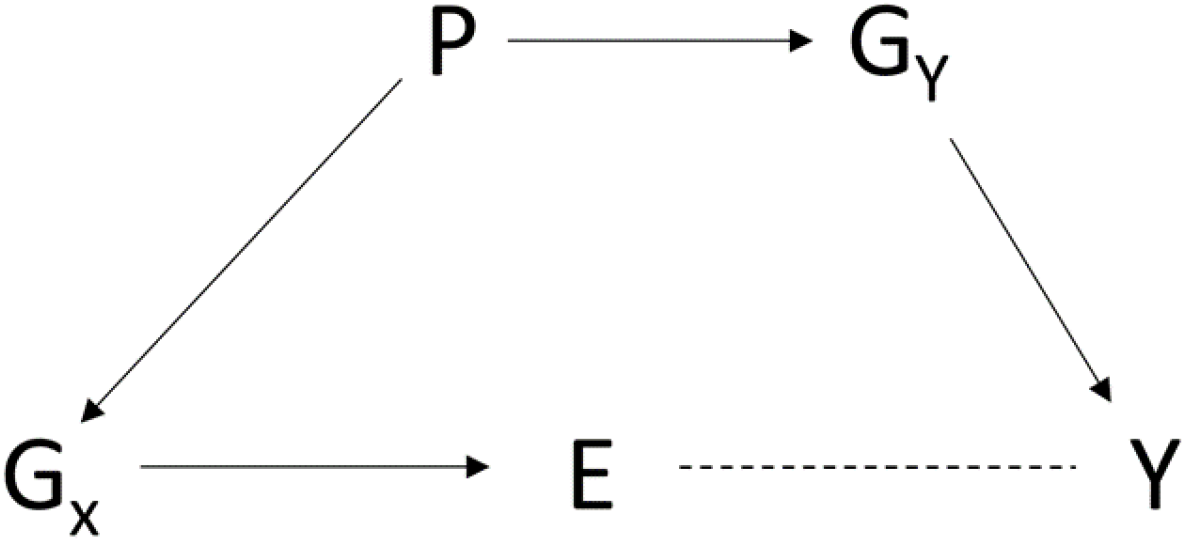
Confounding of the genetic instrument and outcome introduced by population stratification. X is the exposure of interest, Y is the outcome of interest, G_X_ are the genetic variants associated with X used as instruments, P is the population structure, G_Y_ are genetic variants associated with Y through which the population structure affects the outcome. The presence of population stratification creates an association between G_X_ and Y that does not go through X and can induce an apparent association between X and Y even if none exists.

Selection bias, where individuals select to participate in a study, or not, based on their particular phenotypes can also induce bias into any analyses of that study.(14) Particularly, selection bias can induce bias in the associations observed between phenotypes selected on and the genetic variants associated with those phenotypes.(15, 16) The degree to which selection bias is likely to be an issue in any particular study will depend on how that study was recruited. It has been shown in UK Biobank that the characteristics of the study population differ notably from the whole population from which the sample was drawn.(17) Selection bias is a form of collider bias which occurs when the variables of interest independently affect a third variable and so conditioning on this third collider variable will induce an association between the variables of interest.(14, 18) In this case the third variable is participation in the study and conditioning on it is unavoidable as data are only available for the participants. This association could act in either direction and so could either amplify or mask the true association between the variables of interest. (14, 15) Bias of this type can occur even when the genetic instruments are valid instruments for the exposure if the exposure partially determines selection into the study, i.e. selection is dependent on the variables included in the MR estimation.(16) Study participation has shown to be heritable and is influenced by a number of different traits.(19) It is therefore possible for many analyses that participation could be influenced by the traits included in a MR estimation. This setup is illustrated in Fig. 3.

**Figure 3:**
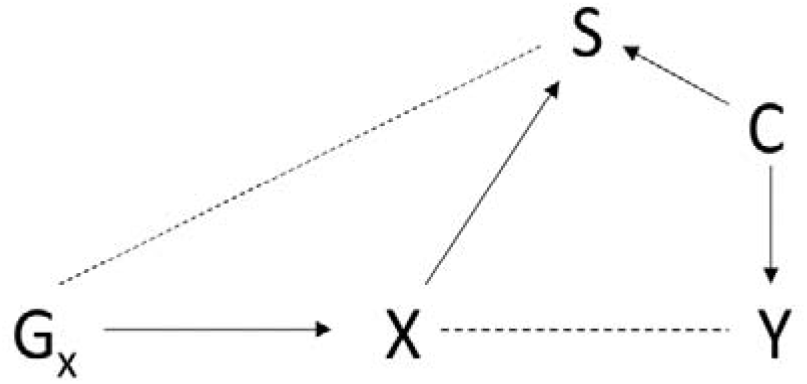
Bias in MR estimates due to selection bias. X is the exposure of interest, Y is the outcome of interest, G_X_ are the genetic variants associated with X used as instruments, S represents selection into the study and C is a cause of the outcome that influences selection into the study. If both the exposure and the cause of the outcome affect selection into the study then conditioning on participation in the study biases estimates of the association between the exposure and the outcome. This will also be true if the outcome influences selection into the study directly.

A method that is often used in observational studies to detect confounding and help assessment of whether a causal relationship exists between an exposure and an outcome is negative control outcome analysis.(20–22) “Negative controls” renames the specificity of associations which Hill considered a factor that should be weighed up in evaluating plausibility of causation in epidemiological studies.(23) Given the dismissal of this criteria in some influential epidemiological texts their rechristening as “negative controls” allowed their relegitimation.(24) Negative control outcome studies compare the association observed between an exposure and an outcome to the association observed between that exposure and a negative control outcome. The negative control outcome variable is chosen to be a variable that is not expected to be associated with the exposure of interest but is expected to be subject to the same unobserved confounding as the exposure and outcome of interest. It follows that if the assumptions hold any association observed between the exposure and the negative control outcome will be due to confounding in the model. Negative control outcome studies have previously also been proposed to detect selection bias in observational studies.(22) We advance the use of negative control outcomes to identify when exposure and outcome phenotypes in an MR analysis may be subject to population stratification or selection bias that has not been fully accounted for in the GWAS and so may induce instrument-outcome confounding or mis-estimation of the SNP exposure relationship and consequently bias the results obtained.

Negative control outcome methods applied to an MR analysis have been used in a limited number of studies previously to detect potential pleiotropy and provide additional evidence on the validity of the MR study.(25–27) These negative controls where however not used in the detection of potential population stratification or selection bias. We illustrate this method through estimation of the potential for population stratification in 33 preselected exposure–indexing GWAS included in MR Base.(28) We detect potential population stratification by estimating the effect of each phenotype on self-reported tanning ability and self-reported natural hair colour, variables that are likely to be highly affected by population stratification but that are largely determined at birth and are not expected to be truly affected by any of the phenotypes considered. Our results from this study show that the GWAS of adiposity related phenotypes and education are likely to be affected by population stratification and/or selection bias. Any MR involving these phenotypes is therefore potentially subject to bias.

## Method

Two-sample summary data MR compares the association of a set of SNPs with the exposure and outcome to determine the effect of the exposure on the outcome. It is explained in detail elsewhere.(6, 7) We propose running two additional MR sensitivity analyses for any MR study where population stratification or selection bias is thought to potentially affect the result.

1. An MR analysis to estimate the effect of the exposure of interest on the negative control outcome.
2. An MR analysis to estimate the effect of the outcome of interest on the negative control outcome (i.e. the outcome of interest becomes the exposure in this analysis).

Any effects detected in these analyses would indicate the potential presence of population stratification or selection bias in the GWAS of the phenotype of interest and therefore possible bias in an MR analysis including that phenotype. The negative control outcomes should be selected based on the same criteria that have been traditionally used in epidemiological studies; i.e. they should not be expected to be dependent on the phenotypes of interest in the analysis but should be affected by the same confounding. In order to satisfy the assumption that the negative control outcome is not actually caused by the exposure we propose using phenotypes that are determined before the exposure and the outcome in the negative control MR study. The phenotype for the negative control outcome should also be selected to be thought to be affected by the population stratification or selection bias. For bias caused by population stratification such variables could include; hair colour, eye colour or skin tone. For bias caused by selection into a study such variables could include educational attainment, which is usually determined in early adulthood and influences participation, for later life exposures or very early life variables such as place of birth. If there is no instrument-outcome confounding this analysis will give a null result. As the negative control outcomes are largely predetermined relative to the exposure and outcome and so cannot depend on either any association of the SNPs with the negative control outcome must be driven by some other mechanism. This could take the form of pleiotropy due to the SNPs having an effect either directly on the negative control outcome or on another phenotype that then affected the negative control outcome, illustrated in Fig. 4. However if this pleiotropy only affects some SNPs it would be detected by conventional pleiotropy robust estimation methods (29–31). Alternatively, the observed effect of the phenotype on the negative control outcome could be due to instrument-outcome confounding. In this case conventional pleiotropy robust methods would not detect an effect as the confounding would affect all of the SNPs. Evidence of an effect of the exposure and outcome on the negative control outcome indicates that an MR study of the exposure on the outcome is also likely to be biased.

**Figure 4:**
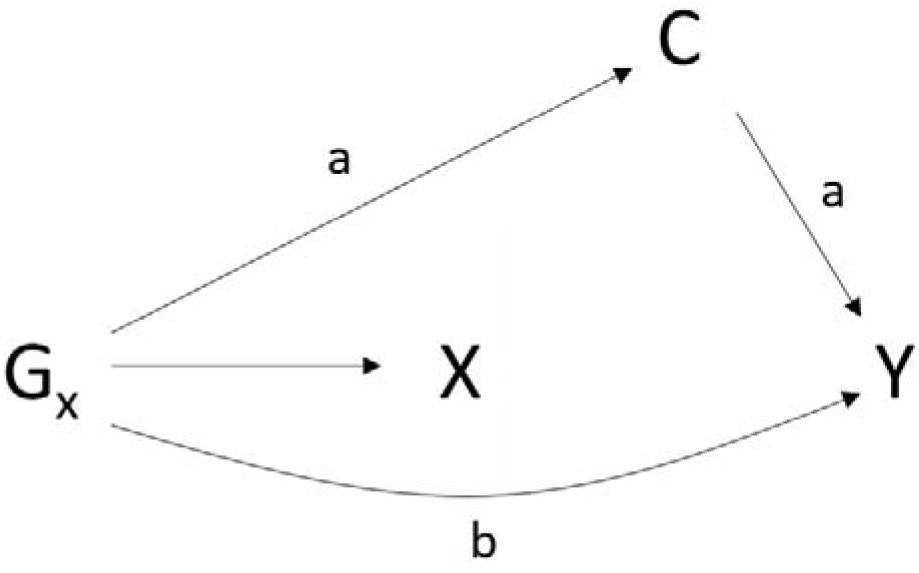
Mechanisms through which pleiotropy can cause bias in MR estimates. X is the exposure of interest, Y is the outcome of interest, G_X_ are the genetic variants associated with X used as instruments, C is an unobserved variable that has a causal effect on Y. Pleiotropy will cause bias in MR estimates if either both edges marked a or the edge marked b are present. Pleiotropy in MR studies is explained in detail elsewhere (3).

## Application

To illustrate the use of negative control outcomes in MR studies we investigated the effect of a range of exposures on self-reported tanning ability and natural hair colour from UK Biobank as a negative control outcome in two-sample summary data MR to detect population stratification in these exposures. Between 2006 to 2010 the UK Biobank study enrolled 500,000 individuals aged between 40 and 69 at baseline across 22 assessments centres in the United Kingdom.(32) Data were collected based on clinical examinations, assays of biological samples, detailed information regarding self-reported health characteristics and genome-wide genotyping.(33) In total, 12,370,749 genetic variants in up to 463,005 individuals were available for analysis as described previously.(34) UK Biobank received ethical approval from the Research Ethics Committee (REC reference for UK Biobank is 11/NW/0382).

For their tanning response to sun exposures individuals were asked “What would happen to your skin if it was repeatedly exposed to bright sunlight without any protection?” with four potential responses which ranged from get very tanned (given a score of 1) to never tan and always burn (given a score of 4). A higher score is therefore associated with fairer skin that is less prone to tanning. A GWAS of this question was conducted by the MRC IEU (34) and included in MR Base.(28) For natural hair colour individuals were asked “What best describes your natural hair colour? (If your hair colour is grey, the colour before you went grey)” with 5 valid potential responses; Blonde, Red, Light brown, Dark brown or Black. We categorised these responses as 1. blonde, 2. red, 3. light brown 4. dark brown and 5. black in accordance with a previous GWAS of hair colour which included UK Biobank (35). The association between genetic variants and outcomes in the UK Biobank study were assessed using the software BOLT-LMM (34, 36). This approach applies a Bayesian linear mixed model to evaluate the association between each genetic variant across the human genome in turn with the analysed outcome accounting for both relatedness and population stratification.(11) Age at baseline, sex and type of genotyping array were added as covariates in the model. As tanning ability and hair colour are largely determined at birth and are is highly dependent on variations in an individual’s ancestral background they should not depend on exposures experienced during an individual’s lifetime.

We preselected 50 characteristics or risk factors with GWAS data available in MR base as our example phenotypes. These phenotypes were all selected to have male and female participants from a mixed or European population that did not include UK Biobank. Where multiple GWAS for the same phenotype where available we chose only the most recent relevant one available in MR base at the time of analysis, however we retained in the analysis similar (but not exactly equivalent) phenotypes such as BMI and Waist to hip ratio. We excluded GWAS that included UK Biobank to avoid the potential for winner’s curse from selecting the exposure and the outcome from the same sample. A full list of the phenotypes included in the analysis are given in Table S1. From these preselected phenotypes we excluded one GWAS due to the information available in MR base not matching that given in the paper and 16 with fewer than 5 genome-wide significant SNPs available as instruments leaving us with 33 phenotypes for analysis.

For each of our 33 exposures we calculated the IVW effect for that exposure on tanning ability and hair colour. For those exposures which showed evidence of an effect on each negative control outcome we also report the MR Egger(31), weighted mode(30) and weighted median(29) effects as sensitivity analyses. Weighted mode and median estimates detect if the association observed is driven by outlying SNPs. However, if population stratification or selection bias is driving the results seen this would not be expected to be due to an effect of a small number of outlying SNPs but due to an effect across all of the SNPs used as instruments. We would therefore still expect to estimate an effect of the trait on the negative control outcome in each case. MR Egger accounts for violation of IV assumptions 2 and 3 that satisfy the InSIDE assumption. This assumption states that the bias on the outcome is independent of the strength of the SNP on the exposure. Bias due to population stratification or selection bias may satisfy this assumption if it applies equally across the SNPs. However, MR Egger has low power to detect effect estimates and so it is often not possible to determine whether the lack of an association in an MR Egger estimation that was observed in an IVW analysis is due to bias in the IVW estimation or low power in the MR Egger estimates. All analyses were conducted using the package “TwoSampleMR” in R.(28)

Estimated effects of the genetic liability towards each exposure on tanning ability from our IVW analysis are given in Fig. 5, with full details given in Supplementary Table S1. These results show that a number of the exposures considered appear to have a causal effect of genetic liability towards that exposure on tanning ability. Table 1 gives the estimated effect sizes for all results with a p-value of less than 0.05 in the IVW analyses. Although we have conducted multiple tests in this analysis as many of the phenotypes we consider are related these tests are not independent. We therefore suggest here that this gives a potential indication of whether results warrant further investigation for potential bias rather than a hard cut off for whether these results are of interest. The traits with an effect on tanning fall into three categories; adiposity related traits, bowel disease and years of schooling. The majority of the GWAS studies included adjusted for population stratification (using principal components or alternative methods) suggesting that this adjustment alone is not sufficient to remove all structural bias in the data. The MR Egger results suffer from high levels of uncertainty due to low power but estimated the same direction of effect in all but three of these exposures. The weighted mode and weighted median estimates supported the overall results with all of the results showing the same direction of effect as the IVW results and only 3 of the 12 results not replicating in at least one of the weighted mode or weighted median estimates.

**Table 1:** Full MR results for exposures which show potential association with tanning ability. Results from IVW, MR Egger, Weighted Mode and Weighted Median analyses for those phenotypes which indicated a potential effect on tanning ability from an MR analysis of 33 preselected phenotypes on tanning ability. P-values in parentheses

| **Exposure** | **No. SNPs** | **IVW** | **MR Egger** | **Weighted Mode** | **Weighted Median** |
| --- | --- | --- | --- | --- | --- |
| *Celiac disease(40)* | 13 | 0.007<br>(5.29E-05) | 0.007<br>(0.033) | 0.007<br>(0.006) | 0.006<br>(0.003) |
| *Triglycerides(41)* | 54 | 0.042<br>(8.46E-04) | 0.027<br>(0.202) | 0.011<br>(0.437) | 0.016<br>(0.166) |
| *Inflammatory bowel disease(42)* | 63 | 0.009<br>(1.13E-03) | 0.009<br>(0.203) | 0.003<br>(0.560) | 0.004<br>(0.186) |
| *Body mass index(43)* | 79 | 0.049<br>(2.05E-03) | 0.055<br>(0.163) | 0.040<br>(0.036) | 0.049<br>(0.005) |
| *Years of schooling(44)* | 70 | 0.074<br>(7.71E-03) | 0.051<br>(0.724) | 0.103<br>(0.034) | 0.092<br>(1.04E-04) |
| *Overweight(45)* | 14 | 0.036<br>(0.018) | -0.025<br>(0.630) | 0.029<br>(0.032) | 0.028<br>(0.014) |
| *Extreme height(45)* | 45 | -0.013<br>(0.025) | -0.043<br>(0.118) | 0.000<br>(0.928) | -0.003<br>(0.240) |
| *HDL cholesterol(41)* | 87 | -0.034<br>(0.029) | -0.008<br>(0.783) | -0.019<br>(0.019) | -0.021<br>(0.033) |
| *Obesity class 2(45)* | 11 | 0.011<br>(0.030) | 0.002<br>(0.927) | 0.011<br>(0.129) | 0.012<br>(0.022) |
| *Waist circumference(46)* | 46 | 0.050<br>(0.030) | 0.097<br>(0.119) | 0.038<br>(0.110) | 0.054<br>(0.009) |
| *Obesity class 1(45)* | 17 | 0.020<br>(0.038) | -0.009<br>(0.735) | 0.013<br>(0.133) | 0.014<br>(0.045) |
| *Childhood obesity(47)* | 5 | 0.028<br>(0.040) | -0.088<br>(0.444) | 0.015<br>(0.119) | 0.018<br>(0.009) |

**Figure 5:**
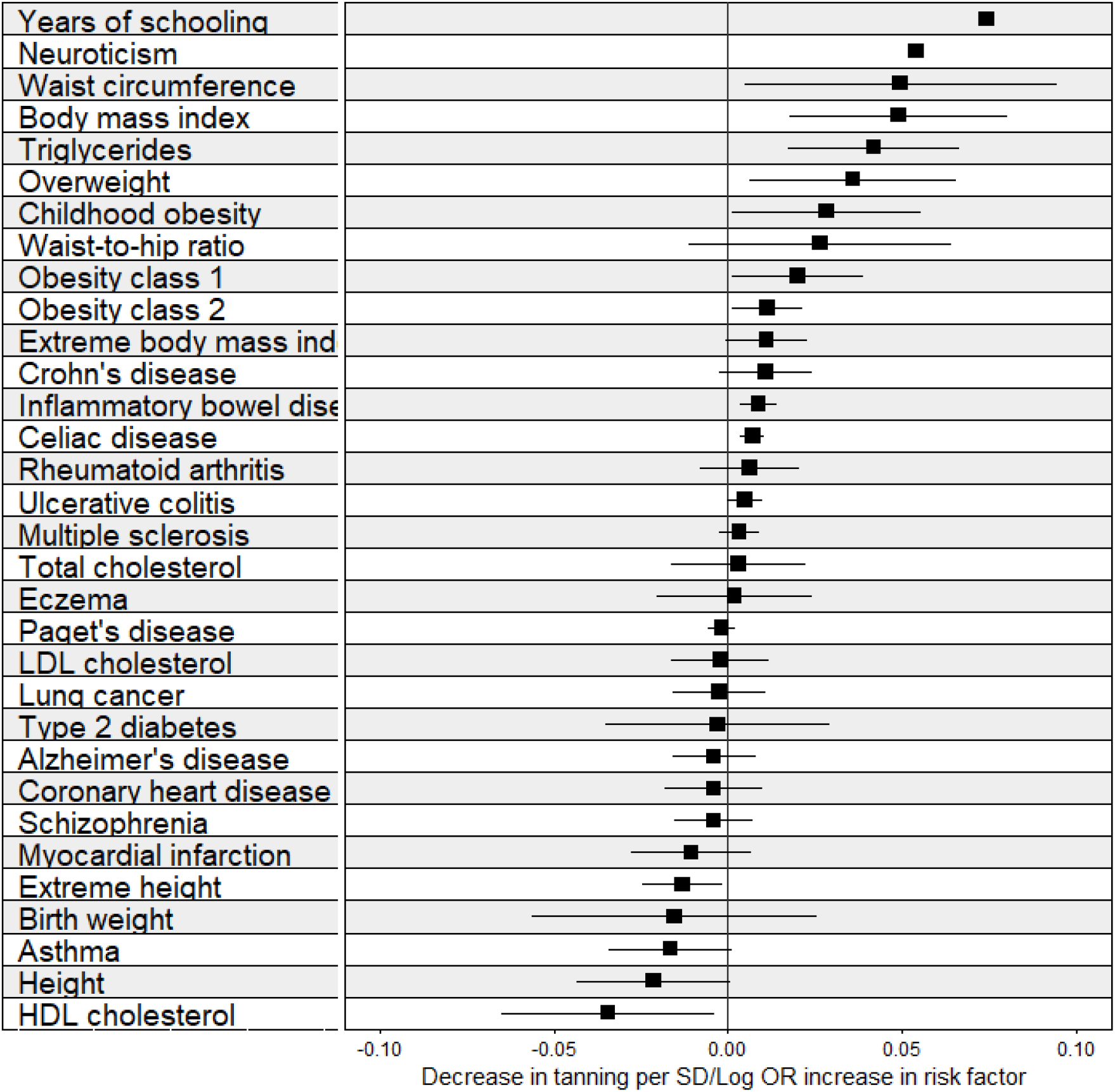
IVW estimates from MR analyses on self-reported tanning ability. IVW results from MR analyses of 33 preselected traits on tanning ability. A higher score indicates less being less likely to tan and more likely to burn when exposed to strong sunlight. Full results from these analyses are given in Table S1.

Estimated effects of the genetic liability towards each exposure on hair colour from our IVW analysis are given in Fig. 6, with full details given in Supplementary Table S1. Results from the IVW analyses and sensitivity analyses for exposures with a p-value <0.05 (as a heuristic for presentation) in the IVW analysis are given in Table 2. These results show a very similar pattern to the results for tanning ability with celiac disease and adiposity and education related traits showing a potential effect on hair colour. Four traits (Triglycerides, Years of schooling, obesity class 2, and celiac disease) showed evidence of an effect on both tanning ability and hair colour.

**Table 2:** MR results for exposures which show potential association with hair colour. Results from IVW, MR Egger, Weighted Mode and Weighted Median analyses for those phenotypes which indicated a potential effect on tanning ability from an MR analysis of 33 preselected phenotypes on self-reported natural hair colour. P-values in parentheses.

| **Exposure** | **No. SNPs** | **IVW** | **MR Egger** | **Weighted Mode** | **Weighted Median** |
| --- | --- | --- | --- | --- | --- |
| *Triglycerides(41)* | 54 | -0.053<br>(0.0004) | -0.059<br>(0.0217) | -0.037<br>(0.0078) | -0.032<br>(0.0063) |
| *Total Cholesterol(41)* | 87 | -0.028<br>(0.0050) | -0.018<br>(0.2807) | -0.026<br>(0.0005) | -0.026<br>(0.0022) |
| *Years of schooling(44)* | 71 | 0.070<br>(0.0095) | 0.069<br>(0.6241) | 0.112<br>(0.0658) | 0.074<br>(0.0021) |
| *Obesity Class 2(45)* | 11 | 0.010<br>(0.0097) | -0.011<br>(0.3533) | 0.002<br>(0.7136) | 0.004<br>(0.4237) |
| *Extreme BMI(45)* | 7 | 0.008<br>(0.0280) | -0.017<br>(0.3745) | 0.001<br>(0.8145) | 0.006<br>(0.1723) |
| *LDL Cholesterol(41)* | 79 | -0.020<br>(0.0304) | -0.016<br>(0.2390) | -0.020<br>(0.0008) | -0.025<br>(0.0011) |
| *Celiac Disease(40)* | 13 | -0.004<br>(0.0395) | -0.007<br>(0.0259) | -0.005<br>(0.0375) | -0.007<br>(0.0030) |

**Figure 1:**
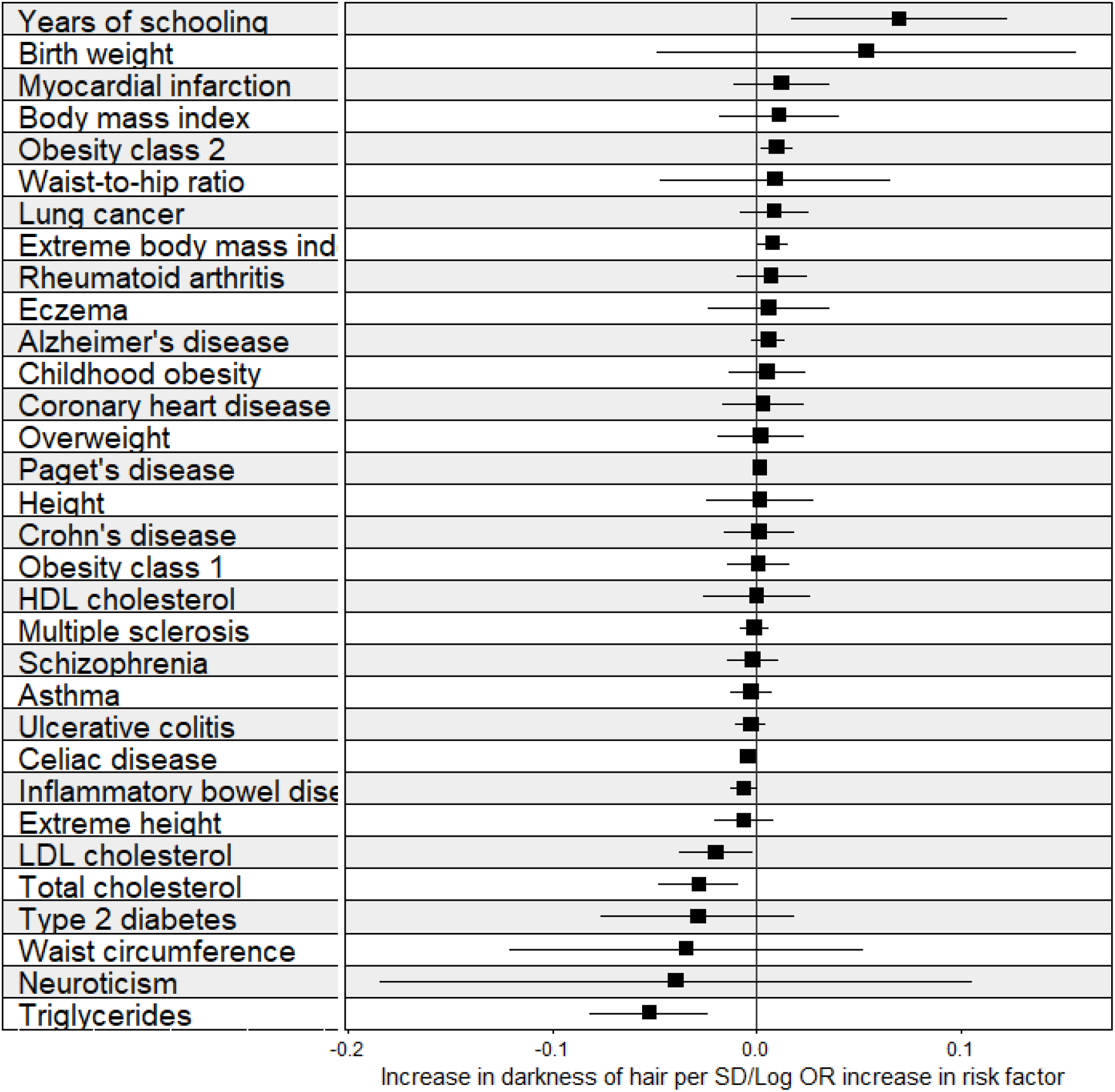
IVW estimates from MR analyses on self-reported natural hair colour. IVW results from MR analyses of 33 preselected traits on hair colour. A higher score indicates darker hair colour. Full results from these analyses are given in Table S1.

## Discussion

In this paper we describe the use of variables that are predetermined relative to the phenotypes of interest but are likely to be subject to population stratification as negative control outcomes within an MR analysis to detect population stratification or selection bias. The method we describe is easy to implement with currently available software and data.

Our applied analysis conducts a negative control outcome MR analysis of 33 preselected phenotypes on tanning ability and natural hair colour to detect the potential for population stratification in the GWAS of these phenotypes. We find a range of phenotypes are potentially affected by population stratification, particularly a number of phenotypes related to BMI, height and educational attainment. Any association between these variables shown by an MR analysis could be due to population stratification introducing the apparent association. This result is supported by a recent study using within-family MR analyses which showed the observed associations from MR analyses between height and education and BMI and education attenuated once family effects where controlled for (13). One key advantage of within-family MR over our method is that is can provide MR causal estimates adjusted for the bias due to population stratification. However, within family MR requires a large sample of related individuals and cannot be conducted with standard GWAS results. The method we propose can detect potential population stratification in samples that do not contain related individuals and using existing summary data.

Genetic liability for celiac disease was associated with both tanning ability and hair colour however this GWAS did not account for population stratification suggesting that the adjustments for population stratification included in GWAS studies do mitigate the effects of population stratification to some degree. However, a number of the other exposures which were found to be associated with tanning ability and/or hair colour did include adjustment for population stratification in the GWAS suggesting that this adjustment does not fully mitigate the effects. Examination of the extent to which adjustment for population stratification through inclusion of principal components or alternative adjustments such as using a BOLT-LMM model (36) in GWAS can mitigate the problems of bias due to instrument-outcome confounding in MR studies is an area for future research.

Although we have focused on the application to two phenotypes that are indicators of population stratification as negative control outcomes this method could equally be applied to detect selection bias. Negative control outcomes to detect selection bias in GWAS results would include phenotypes that are expected to affect participation in a study but that are predetermined relative to the exposure and outcome considered in the MR analysis. Such negative control outcomes could include early life variables such as place of birth or education. Alternatively, participation can be examined directly in birth cohort studies which are followed up over time and GWAS results from these studies could be used as a negative control outcome. (19)

LD score regression is a method that attempts to separate out biological and confounded genetic signals and so can also be applied in an MR setting to determine whether there is a causal (biological) effect of an exposure on an outcome or if an observed association is due to confounding.(37) LD score regression however does not give the appropriate results if the GWAS being considered have been performed using a linear mixed model. Our method therefore provides a complimentary approach to LD score regression that does not depend on the method used to estimate GWAS associations. Additionally, LD score regression incorporates data from the entire genome whereas the used of negative controls outcomes proposed here only uses SNPs strongly associated with the phenotypes of interest. This is potentially more relevant to bias in MR analyses which uses SNPs associated with the exposure to estimate the causal effect of the exposure on the outcome.

There are a number of weaknesses with our method that should be considered. This method is only able to detect bias as far as it affects the chosen negative control outcome and therefore no detected effect of the phenotype on the negative control outcome does not mean that the phenotype, and any associated MR analysis, is necessarily free from bias. This limitation can be mitigated by choosing negative control outcomes that are likely to be highly population stratified or selected on as far as they are available.

Negative control outcome calibration has been proposed for observational negative control studies to adjust the effect of exposure of interest on the outcome for the bias detected by the negative control outcome.(38) We believe that mechanical application of such an approach should be avoided due to the strong assumptions required for such calibration to give reliable estimates.(39) A key assumption for such an approach to work is that the model fully identifies the effect of the exposure on the outcome and negative control outcome such that the size of the effect of the bias on the outcome can be determined once and differences in scale of the outcome and negative control have been taken into account. In the context of MR this is not a reasonable assumption as this assumption would require the instrument-outcome confounding to have exactly the same effect in the exposure, outcome and negative control outcome and so if bias is detected this method does provide a method to correct the estimated effect. However, the size and direction of the estimated effect on the negative control outcome could be used as an indicator for a sensitivity analysis which considered whether bias of up to, for example, 5 times that estimated by the negative control outcome would change the conclusions from the main MR analyses.

An extension to this method is to consider the use of similar phenotypes considered as negative control outcomes here as negative control exposures. Such an approach provides an obvious complement to the approach considered here, however, the assumptions required and implications of such an analysis are notably different to those for a negative control outcome study and therefore we leave this as an area for future research.

## Data and Code Availability

Data for all outcomes considered and ‘tanning response to sun exposure’ exposure are available as part of the R package “TwoSampleMR”. Code for the negative control analyses conducted are available at https://github.com/eleanorsanderson/MR-negativecontrols. The GWAS of natural hair colour was conducted using data from UK Biobank (https://www.ukbiobank.ac.uk/) under application number 15825.

## Acknowledgements

ES and GDS work in a unit supported by the MRC (MC_UU_00011/1), TGR is a UKRI Innovation Research Fellow (MR/S003886/1).

## Author Contributions

ES and GDS conceived the study. TGR conducted the GWAS of hair colour. ES conducted all other analyses and wrote the first draft of the manuscript. All authors edited and approved the manuscript.

