## Supplementary material for "The use of negative control outcomes in Mendelian Randomisation to detect potential population stratification or selection bias"

**Table S1 –** All phenotypes included in the estimation and their estimated effect on tanning response and hair colour from a two-sample IVW estimation.

|  | **Skin tone** | | | **Hair colour** | | |  |  |  |  |  |
| --- | --- | --- | --- | --- | --- | --- | --- | --- | --- | --- | --- |
| **Exposure** | **Effect** | **Std. Error** | **P value** | **Effect** | **Std. Error** | **P value** | **No. SNPs** | **Population** | **Sample size** | **Units** | **Year** |
| Alzheimer's disease(1) | -0.004 | 0.006 | 0.5171 | 0.006 | 0.004 | 0.1582 | 20 | European | 74046 | log odds | 2013 |
| Anorexia nervosa(2) |  |  |  |  |  |  | 1 | European | 17767 | log odds | 2014 |
| Asthma(3) | -0.016 | 0.009 | 0.0711 | -0.002 | 0.005 | 0.6319 | 8 | European | 26475 | log odds | 2007 |
| Autism(4)^[1]^ |  |  |  |  |  |  |  | European |  | log odds | 2015 |
| Bipolar disorder(5) |  |  |  |  |  |  | 4 | European | 16731 | log odds | 2011 |
| Birth length(6) |  |  |  |  |  |  | 2 | European | 28459 | SD (cm) | 2015 |
| Birth weight(7) | -0.015 | 0.021 | 0.4658 | 0.054 | 0.052 | 0.3058 | 52 | Mixed | 153781 | SD | 2016 |
| Body Mass Index(8) | 0.049 | 0.016 | 0.0021 | 0.011 | 0.015 | 0.4514 | 79 | Mixed | 339224 | SD (kg/m^2) | 2015 |
| Cardioembolic stroke(9) |  |  |  |  |  |  | 2 | Mixed | 21185 | NA | 2016 |
| Celiac disease(10) | 0.007 | 0.002 | 0.0001 | -0.004 | 0.002 | 0.0395 | 13 | European | 24269 | log odds | 2011 |
| Childhood obesity(11) | 0.028 | 0.014 | 0.0400 | 0.005 | 0.010 | 0.5942 | 5 | European | 13848 | log odds | 2012 |
| Chronic kidney disease(12) |  |  |  |  |  |  | 4 | Mixed | 117165 | log odds | 2015 |
| College completion(13) |  |  |  |  |  |  | 3 | European | 95427 | log odds | 2013 |
| Coronary heart disease(14) | -0.004 | 0.007 | 0.5653 | 0.003 | 0.010 | 0.7512 | 39 | Mixed | 184305 | log odds | 2015 |
| Crohn's disease(15) | 0.011 | 0.007 | 0.1109 | 0.001 | 0.009 | 0.8884 | 122 | European | 51874 | log odds | 2015 |
| Depressive symptoms(16) |  |  |  |  |  |  | 1 | European | 161460 | SD | 2016 |
| Difference in height between adolescence and adulthood(17) |  |  |  |  |  |  | 1 | European | 9228 | SD | 2013 |
| Difference in height between childhood and adulthood(17) |  |  |  |  |  |  | 1 | European | 10799 | SD | 2013 |
| Eczema(18) | 0.002 | 0.011 | 0.8663 | 0.006 | 0.015 | 0.6908 | 12 | European | 40835 | log odds | 2014 |
| Extreme body mass index(19) | 0.011 | 0.006 | 0.0636 | 0.008 | 0.004 | 0.0280 | 7 | European | 16068 | log odds | 2013 |
| Extreme height(19) | -0.013 | 0.006 | 0.0249 | -0.006 | 0.007 | 0.4119 | 45 | European | 16196 | log odds | 2013 |
| Extreme waist-to-hip ratio(19) |  |  |  |  |  |  | 2 | European | 10255 | log odds | 2013 |
| Gout(20) |  |  |  |  |  |  | 2 | European | 69374 | log odds | 2013 |
| HDL cholesterol(21) | -0.034 | 0.016 | 0.0286 | 0.000 | 0.013 | 0.9858 | 87 | Mixed | 187167 | SD (mg/dL) | 2013 |
| Height(22) | -0.021 | 0.011 | 0.0591 | 0.002 | 0.013 | 0.8919 | 367 | European | 253288 | SD (m) | 2014 |
| Inflammatory bowel disease(15) | 0.009 | 0.003 | 0.0011 | -0.006 | 0.003 | 0.0794 | 63 | European | 34652 | log odds | 2015 |
| Ischaemic stroke(23) |  |  |  |  |  |  | 1 | European | 517 | log odds | 2007 |
| LDL cholesterol(21) | -0.002 | 0.007 | 0.7736 | -0.020 | 0.009 | 0.0304 | 79 | Mixed | 173082 | SD (mg/dL) | 2013 |
| Lung adenocarcinoma(24) |  |  |  |  |  |  | 2 | European | 18336 | log odds | 2014 |
| Lung cancer(24) | -0.002 | 0.007 | 0.7215 | 0.009 | 0.008 | 0.2985 | 5 | European | 27209 | log odds | 2014 |
| Multiple sclerosis(25) | 0.003 | 0.003 | 0.2420 | -0.001 | 0.004 | 0.7630 | 46 | European | 38589 | log odds | 2013 |
| Myocardial infarction(14) | -0.010 | 0.009 | 0.2373 | 0.012 | 0.012 | 0.3135 | 25 | Mixed | 171875 | log odds | 2015 |
| Neuroticism(16) | 0.054 | 0.066 | 0.4100 | -0.040 | 0.074 | 0.5921 | 9 | European | 170911 | SD | 2016 |
| Obesity class 1(19) | 0.020 | 0.010 | 0.0380 | 0.001 | 0.008 | 0.9105 | 17 | European | 98697 | log odds | 2013 |
| Obesity class 2(19) | 0.011 | 0.005 | 0.0297 | 0.010 | 0.004 | 0.0097 | 11 | European | 72546 | log odds | 2013 |
| Obesity class 3(19) |  |  |  |  |  |  | 2 | European | 50364 | log odds | 2013 |
| Overweight(19) | 0.036 | 0.015 | 0.0181 | 0.002 | 0.011 | 0.8450 | 14 | European | 158855 | log odds | 2013 |
| Paget's disease(26) | -0.002 | 0.002 | 0.3849 | 0.002 | 0.002 | 0.3074 | 5 | European | 3440 | log odds | 2011 |
| Parkinson's disease(27) |  |  |  |  |  |  | 4 | European | 1672 | log odds | 2011 |
| Rheumatoid arthritis(28) | 0.006 | 0.007 | 0.3881 | 0.007 | 0.009 | 0.4074 | 47 | Mixed | 80799 | log odds | 2014 |
| Schizophrenia(29) | -0.004 | 0.006 | 0.4838 | -0.002 | 0.006 | 0.7681 | 71 | Mixed | 82315 | log odds | 2014 |
| Squamous cell lung cancer(24) |  |  |  |  |  |  | 4 | European | 18313 | log odds | 2014 |
| Subjective well being(16) |  |  |  |  |  |  | 1 | European | 298420 | SD | 2016 |
| Total cholesterol(21) | 0.003 | 0.010 | 0.7604 | -0.028 | 0.010 | 0.0050 | 87 | Mixed | 187365 | SD (mg/dL) | 2013 |
| Triglycerides(21) | 0.042 | 0.013 | 0.0008 | -0.053 | 0.015 | 0.0004 | 54 | Mixed | 177861 | SD (mg/dL) | 2013 |
| Type 2 diabetes(30) | -0.003 | 0.016 | 0.8646 | -0.029 | 0.024 | 0.2346 | 36 | European | 69033 | log odds | 2012 |
| Ulcerative colitis(15) | 0.005 | 0.003 | 0.0649 | -0.003 | 0.004 | 0.4628 | 85 | European | 47745 | log odds | 2015 |
| Waist circumference(31) | 0.050 | 0.023 | 0.0302 | -0.035 | 0.044 | 0.4334 | 46 | Mixed | 224459 | SD (cm) | 2015 |
| Waist-to-hip ratio(31) | 0.026 | 0.019 | 0.1677 | 0.009 | 0.029 | 0.7500 | 30 | Mixed | 224459 | SD | 2015 |
| Years of schooling(32) | 0.074 | 0.028 | 0.0077 | 0.070 | 0.027 | 0.0095 | 70 | European | 293723 | SD (years) | 2016 |

1. This study was dropped because the description data given in MR base for sample size did not match that given in the paper for this trait.
